# Prior hypoxia exposure enhances murine microglial inflammatory gene expression *in vitro* without concomitant H3K4me3 enrichment

**DOI:** 10.1101/2020.02.03.933028

**Authors:** Elizabeth A. Kiernan, Andrea C. Ewald, Jonathan N. Ouellette, Tao Wang, Avtar Roopra, Jyoti J. Watters

## Abstract

Hypoxia is a component of multiple disorders, including stroke and sleep-disordered breathing, that often precede or are comorbid with neurodegenerative diseases. However, little is known about how hypoxia affects the ability of microglia, resident CNS macrophages, to respond to subsequent inflammatory challenges that are often present during neurodegenerative processes. We therefore tested the hypothesis that hypoxia would enhance or “prime” microglial pro-inflammatory gene expression in response to a later inflammatory challenge without programmatically increasing basal levels of pro-inflammatory cytokine expression. To test this, we pre-exposed immortalized N9 and primary microglia to hypoxia (1% O2) for 16 hrs and then challenged them with pro-inflammatory lipopolysaccharide (LPS) either immediately or 3-6 days following hypoxic exposure. We used RNA sequencing coupled with chromatin immunoprecipitation sequencing to analyze primed microglial inflammatory gene expression and modifications to histone H3 lysine 4 trimethylation (H3K4me3) at the promoters of primed genes. We found that microglia exhibited enhanced responses to LPS 3 days and 6 days post-hypoxia. Surprisingly however, the majority of primed genes were not enriched for H3K4me3 acutely following hypoxia exposure. Using the bioinformatics tool MAGICTRICKS and reversible pharmacological inhibition, we found that primed genes required the transcriptional activities of NF-ĸB. These findings provide evidence that hypoxia pre-exposure could lead to persistent and aberrant inflammatory responses in the context of CNS disorders.

## INTRODUCTION

Hypoxia (Hx) is a component of multiple disorders affecting the central nervous system (CNS), including stroke, cancer, sleep-disordered breathing, and apneas of prematurity (1–5). The most common clinical manifestation of pathological Hx is sleep-disordered breathing, a disorder present in 30% of men and 17% of women aged 30-70, characterized by recurrent bouts of Hx during sleep (6). This disorder causes long-term cognitive deficits (7–11) and increases susceptibility to multiple neurodegenerative diseases (12–16). Despite the common co-occurrence of Hx with neural disease, few studies have investigated how pre-exposure to Hx changes the long-term ability of resident CNS macrophages/microglia to respond to a subsequent inflammatory insult, such as pathogen exposure or a sterile inflammatory stimulus.

In peripheral and CNS immune cells, there is significant cross-talk between the pathways mediating responses to Hx and inflammation (reviewed in (17, 18)), including shared activation of the transcription factor HIF-1α (19), suggesting a mechanism whereby Hx could have long-term effects on microglial inflammatory responses. In normoxia (Nx), pathogen exposure activates HIF-1α and shifts peripheral macrophage metabolism from oxidative phosphorylation to glycolysis, promoting a more pro-inflammatory M1 phenotype (20–22). This metabolic shift “primes” peripheral macrophages and enhances their pro-inflammatory cytokine expression upon subsequent pathogen exposure, even long after the initial pathogen stimulus has been removed (20–22). Although the ability of Hx-induced HIF-1α to similarly prime macrophage/microglia inflammatory responses long-term has not been tested, studies simultaneously exposing macrophages to acute Hx and pro-inflammatory stimuli show that Hx enhances pro-inflammatory gene expression (23) and phagocytic efficiency (24). However, these simultaneous exposures do not elucidate the effects of Hx pre-exposure in the absence of other stimuli on macrophage priming, nor the underlying molecular mechanisms contributing to long-term effects.

Both Hx and macrophage pathogen priming shift cell metabolism to glycolysis (25–27), impacting the availability of cofactors necessary for histone modifications that regulate inflammatory gene programs (21, 28, 29). This suggests that an epigenetic mechanism may underlie long-term pathogen and Hx priming of macrophage inflammatory gene expression. Supporting this idea, macrophages/monocytes primed with diverse stimuli, including β-glucan, lipopolysaccharide (LPS), or tuberculosis vaccine display enriched histone 3 lysine 4 mono- and trimethylation (H3K4Me1 and H3K4me3 respectively) at both pro-inflammatory and glycolytic genes (20, 21, 30, 31). For β-glucan priming, this increase is attributed to inhibition of KDM5, an H3K4me3 histone demethylase, and restoration of KDM5 activity reverses macrophage priming (29). Studies in non-macrophage cells show that Hx, like pathogen priming, also increases global H3K4me3 at glycolytic and pro-inflammatory genes in a KDM5A-dependent manner (32, 33), and interestingly, these changes are independent of HIF-1α (32). Together, these studies demonstrate that both pathogen priming and Hx lead to similar epigenetic changes at pro-inflammatory genes, although it remains unknown if the Hx-induced H3K4me3 present at pro-inflammatory genes is associated with enhanced expression of those genes upon exposure to subsequent inflammatory stimuli, or if these changes specifically occur in CNS macrophages.

Thus, here we tested the hypothesis that Hx pre-exposure enhances (“primes”) microglial pro-inflammatory gene expression in responses to subsequent inflammatory stimuli both acutely, and days after, the initial Hx stimulus. Further, we examined if Hx-induced priming was associated with H3K4me3 enrichment at primed pro-inflammatory gene promoters. Using RNA sequencing and H3K4me3 chromatin-immunoprecipitation sequencing, we found that exposing microglia to Hx prior to an inflammatory challenge with LPS primes pro-inflammatory cytokine gene expression both acutely and long-term. Interestingly, although Hx alone enriched H3K4me3 at glycolysis- and immune-related genes, it did not increase H3K4me3 at the majority of primed genes, suggesting that alternative molecular mechanisms underlie long-term priming effects. The application MAGICTRICKS identified NF-ĸB as common DNA binding factor regulating genes primed by Hx, and its pharmacological inhibition confirmed an important role for this transcription factor in Hx-induced gene priming.

## MATERIALS AND METHODS

### Cell Culture and Reagents

Immortalized murine N9 microglia (RRID:CVCL_0452) were cultured as previously described (34). Cells were verified to be mycoplasma-free. For qRT-PCR experiments, cells were plated in 24-well plates at 5x10^4^ cells/well. For chromatin immunoprecipitation studies, cells were plated at ∼2x10^6^ cells per 10 cm plate. For Hx treatments, cells were exposed to 1-1.5% O2/5% CO2 (Hx) or 21% O2/5% CO2 (Nx) for 16 hrs after which time, vehicle (HBSS) or 100 ng/mL lipopolysaccharide (LPS *E. coli* O111:B4; 500,000 EU/mg; Sigma-Aldrich; St. Louis, MO) was administered for 3 hrs in Hx or Nx, respectively. This level of hypoxia was sufficient to induce hypoxia-sensitive genes and did not result in significant loss of cell viability (Supplemental Figure 1). For NF-κB inhibition studies, vehicle (DMSO) or 3 µM BOT-64 (Abcam; Cambridge, MA) was added prior to the start of Hx. The dose of BOT-64 was determined by analyzing the concentration necessary to inhibit LPS-induced microglial gene expression (data not shown).

Primary microglia were isolated from C57BL/6J (The Jackson Laboratory, Cat# JAX:000664, RRID:IMSR_JAX:000664) mouse line as previously described (35). Cells were plated at 2.5x10^5^ cells/well. For Hx treatments, cells were exposed to 0.1-0.5% O2/5% CO2 (Hx) or Nx for for 24 hrs. The growth medium was replaced with DMEM supplemented with 2 ng/mL murine M-CSF (R&D Systems; Minneapolis, MN) and cultured in Nx for 3 or 6 days prior to LPS treatment.

### Quantitative RT-PCR

qRT-PCR using power SYBR green (ThermoFisher Scientific) was performed as previously described (36) using an Applied Biosystems 7500 Fast Real Time PCR System. All primers (Table 1) were tested for efficiency using serial dilutions, and results were normalized to 18S RNA levels; data analyses were performed using the standard curve analysis method (37, 38). The 18S ribosomal transcript was used as a house keeping gene as our hypoxia treatments did not alter its expression (Table 2).

**Table 1.**
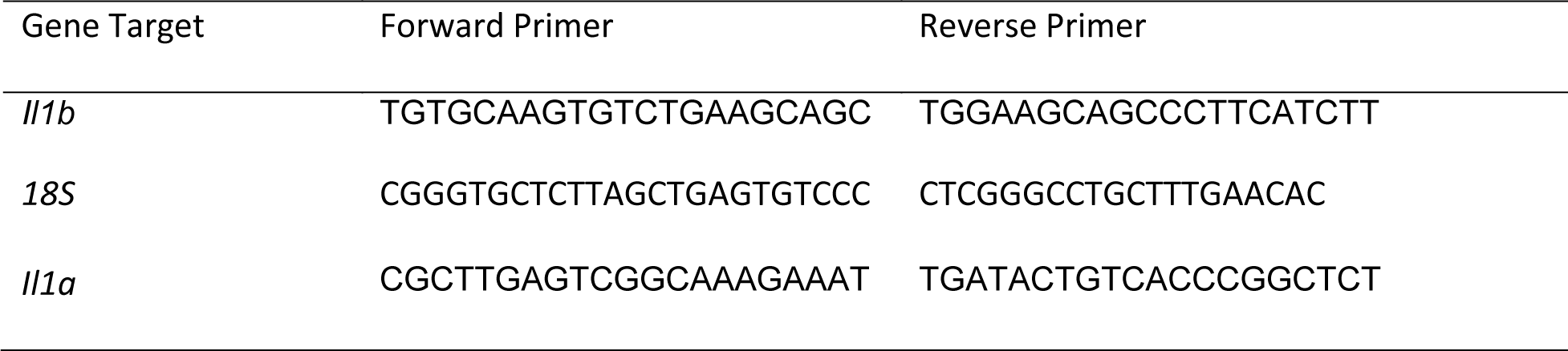
Primers used for quantitative PCR

**Table 2.** Microglial 18S CT values

| Treatment | N9 | Std. Error | N | Primary | Std. Error | N |
| --- | --- | --- | --- | --- | --- | --- |
|  | Mean CT |  |  | Mean CT |  |  |
| Nx | 10.698 | 0.170 | 10 | 10.583 | 0.173 | 8 |
| Nx LPS | 10.708 | 0.196 | 10 | 10.706 | 0.192 | 8 |
| Hx | 10.908 | 0.187 | 10 | 10.765 | 0.178 | 8 |
| Hx LPS | 11.000 | 0.190 | 10 | 10.760 | 0.204 | 8 |

### RNA Sequencing and Analyses

For comparisons between N9s exposed to 16 hrs Nx or Hx, total RNA was isolated using Trizol as previously described (36) and submitted to the UW-Madison Biotechnology Resource Center for sequencing and quality control. Stranded cDNA libraries were prepared using the Truseq Stranded mRNA kit (Illumina; San Diego, CA). Sequencing was performed using an Illumina HiSeq 2000. After sequencing and quality control, fastq files containing ∼ 35 million reads for each biological replicate (n = 3/treatment) were analyzed using the Tuxedo suite (39). Alignment to the mm10 genome was made using Tophat2.1.0 and Bowtie1. Differential analysis was performed using Cufflinks followed by CuffDiff. For comparisons between N9s exposed Nx LPS and Hx LPS, total RNA was isolated using Trizol and samples were submitted to Novogene for library construction and paired-end (PE-150) sequencing with an Illumina NovaSeq. Fastq files containing ∼ 30 million reads for each biological replicate (n = 5/treatment) were similarly aligned using Tuxedo suite with Tophat 2.1.0. Differential gene expression was analyzed with Cuffdiff. Differential expression was considered statistically significant using an FDR<5%. Results were uploaded to NCBI Geo Datasets (GEO GSE108770; https://www.ncbi.nlm.nih.gov/geo/query/acc.cgi?acc=GSE108770).

### MAGICTRICKS Analyses

Putative transcription factors and cofactors driving gene expression changes were identified using MAGICTRICKS (40). To prepare data for these analyses, we first used the Cuffdiff output to identify genes that were significantly upregulated in Hx LPS relative to Nx LPS (FDR<5%). Next, we subtracted those genes upregulated in Hx relative to Nx from this list. This resulted in a list of “primed” genes. A background list of genes was created using all genes that were expressed according to Cuffdiff output comparing Hx LPS vs. Nx LPS (i.e. genes that had a status of “ok”). These two lists (background and primed genes) were formatted according to MAGICTRICKS instructions and run on the MAGICTRICKS software. Cumulative distribution function graphs (CDFs) and summary Scores were generated in the MAGICTRICKS software.

### Chromatin Immunoprecipitation qPCR

N9 microglia were fixed in 1% formaldehyde at room temperature for 2 mins and lysed in 150 mM NaCl, 10% glycerol, and 50 mM Tris (pH 8.0). Chromatin was sonicated to a fragment size of ∼500 bp (assessed by gel electrophoresis). Protein was quantified by BCA Assay (Pierce; Rockford IL). Chromatin (125 µg/IP) was incubated overnight in 2 µg of either rabbit polyclonal anti-histone 3 lysine 4 trimethyl (H3K4me3; Abcam, Cat# ab8580, RRID:AB_306649) or anti-Histone 3 (Abcam, Cat# ab1791, RRID:AB_302613) antibodies, and 10% input was set aside until purification. Simultaneously, protein G Sepharose beads (GE Healthcare) were blocked overnight with 0.5 mg BSA. Cleared Sepharose G beads were added to the sheared chromatin for 1 hr, and the chromatin was washed and reversed cross-linked as previously described by other research groups (41). DNA was purified by phenol chloroform extraction and quantified (primers in Table 1) using an ABI7500 FAST and POWER SYBR Green (ThermoFisher Scientific). H3K4me3 enrichment was quantified relative to the 10% input control.

### H3K4me3 Chromatin Immunoprecipitation Sequencing (ChIP-seq) and Analyses

Chromatin was immunoprecipitated with anti-H3K4me3 antibodies (n=3 independent biological replicates) and submitted to the UW-Madison Biotechnology Center for Illumina sequencing. Fastq files were processed by ProteinCT (Madison, WI) using Bowtie2 to the mm10 genome. Combined tag directories for each treatment (including three 10% inputs samples) were created using HOMER (42). To examine overall H3K4me3 peaks, unique peaks were called using Homer’s findPeaks.pl script using the default four-fold above their respective input controls. Results were uploaded to Geo Datasets (GEO GSE108770; https://www.ncbi.nlm.nih.gov/geo/query/acc.cgi). Tag density files (TDF) were visualized with the Integrative Genomics Viewer (IGV) (43).

### Analysis of H3K4me3 ChIP-seq for beta-glucan primed human monocytes

To analyze previously published data (GEO GSE34260) of human monocytes primed with β-glucan (20), we extracted H3K4me3 ChIP-seq fastq files for replicate human monocyte samples (n = 2) treated with either RMPI or β-glucan. Fastq files were aligned to the hg38 genome with Bowtie2. Tag directories were created using Homer’s makeTagDirectory, and consensus directories for replicates were made using makeTagDirectory with –single parameter. Because this dataset had no input control, the RPMI consensus Tag Directory was used as background for calling peaks with Homer’s findPeaks using the – style factor parameter. Peaks were called when they were 1.4-fold higher than background. To better compare β-glucan-induced peaks to Hx-induced H3K4me3 peaks, we similarly reanalyzed the N9 Nx and Hx H3K4me3 Tag Directories using the consensus Nx H3K4me3 tag directory as background instead of the 10% input. Gene ontology for Biological Process was performed on H3K4me3 peaks that were enriched between -1kb and +300 bp of the transcription start site using STRING analyses (44).

### Gene Ontology Analyses

Gene ontology for KEGG Pathway enrichment and for Biological Process was performed using the STRING software (44). For these analyses, background gene lists were provided based on the Cuffdiff output for RNAseq comparisons. Genes that had a status of “ok” on Cuffdiff output were considered expressed genes and used as background for bioinformatic analyses. For ChIP-seq analyses, whole genome was used as background.

### Flow Cytometry for Cell Viability

Microglia were stained with eFluor 780 live/dead (eBioscience 65-0865-14). Flow cytometry was performed using a BD Fortessa Flow Cytometer (gating strategy, Supplemental Figure 1). FCS files were analyzed using FlowJO v.10.0.7 software.

### Statistics

All statistical analyses for non-genomic work were performed using SigmaPlot (Systat Software, San Jose CA) unless otherwise noted. Tests for normality were done prior to statistical tests that require normality. Non-normal data was log transformed and assessed for a log-normal distribution. The Grubb’s Test was used to identify outliers. Statistical significance (set at p < 0.05) was determined using a Two-way repeated measures ANOVA (Two-way RM ANOVA), or paired t-tests, followed by a Holm-Sidak post hoc test when applicable. P-values between 0.05-0.1 were considered to be statistical trends.

## RESULTS

### Hx induces global gene transcription changes in microglia and upregulates immune-and glycolysis-related genes

To identify the effects of Hx pre-exposure on subsequent microglial responses to an inflammatory challenge, we first tested the global effects of Hx alone on microglial gene transcription. We exposed microglia to 16 hrs of Hx (1% O2) or Nx (room air), and isolated total RNA for sequencing. Differential expression analyses with Cufflinks revealed large changes in the transcriptional program, with Hx upregulating 1,119 genes and downregulating 1,432 genes relative to Nx (Figure 1A). Similar to the effects of Hx reported in other cell types (32), KEGG Pathway analysis demonstrated that genes upregulated by Hx were primarily involved immune system and metabolic function. Specifically, genes upregulated by Hx were significantly enriched in the categories of “cytokine-cytokine receptor interactions”, “glycolysis”, and “HIF-1α signaling pathway” (Figure 1B). Many of the genes enriched in these categories were chemokine receptors, such as the fractalkine receptor “CX3CR1”, or genes associated with glucose metabolism such as phosphofrutokinase Liver Type (PFKL) and Hexokinase 2 (HK2; Figure 1C). Together, these results support that Hx induces a global shift in microglial gene expression, and similar to other models of immune gene priming, that Hx upregulates a transcriptional program related to glucose metabolism.

**Figure 1.**
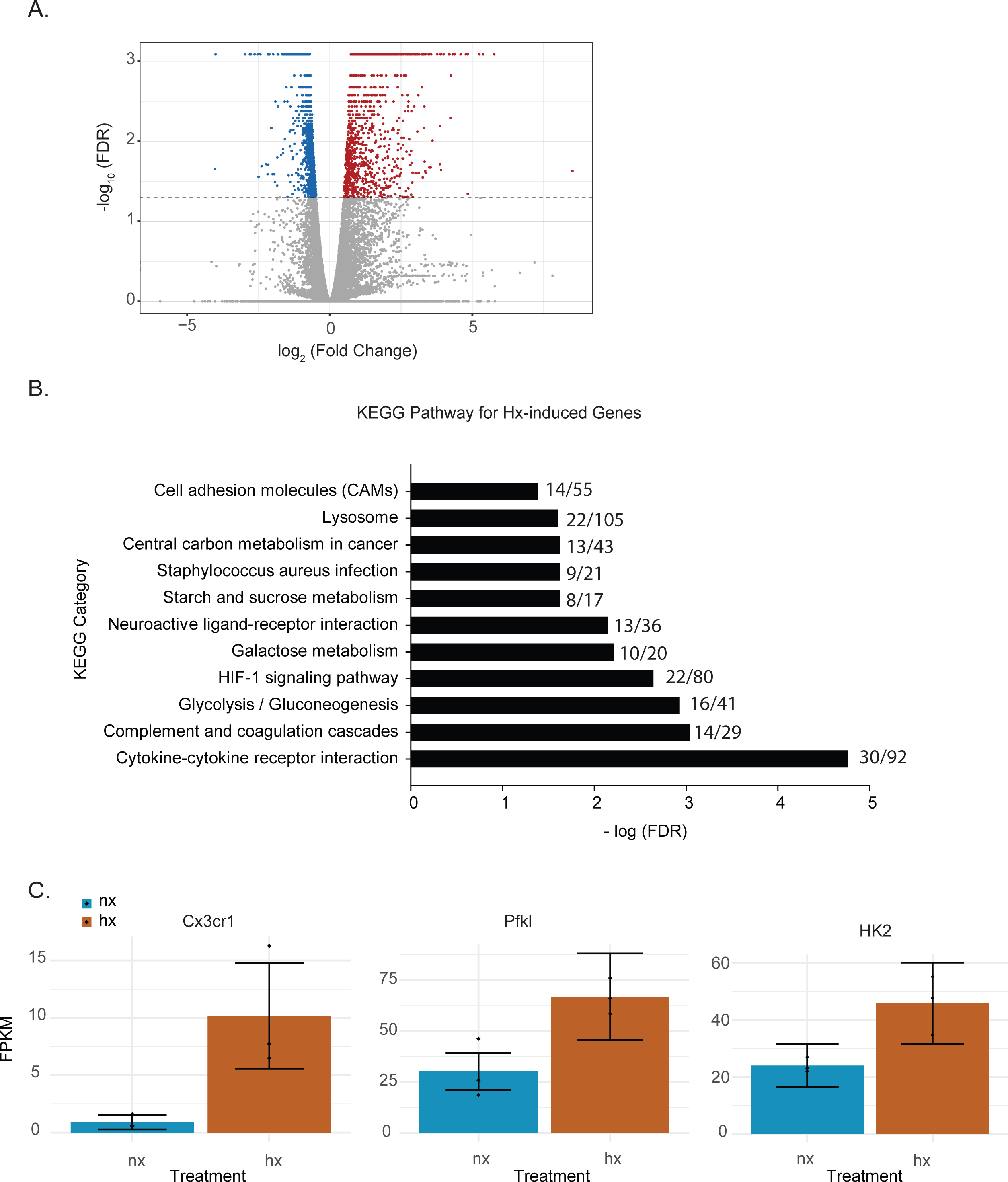
Hypoxia changes microglial gene expression and upregulates metabolic- and cytokine receptor-related genes. A) Volcano plot of microglial differential gene expression after Hx exposure. Red dots are upregulated genes, blue dots are downregulated genes. B) Upregulated genes were analyzed for KEGG Pathway enrichment using STRING software. The numbers at the end of each bar represent the observed/expected genes in each category. C) FPKMs of genes upregulated by Hx that are in the categories of “cytokine-cyotkine receptor interaction” or “HIF-1 Signaling”. Results are derived from n = 3 biological replicates/treatment.

### Hx primes pro-inflammatory gene transcription

We next compared the transcriptomic effects of Hx to the inflammatory stimulus LPS and tested if Hx pre-exposure could enhance LPS-induced gene transcription. We exposed microglia to 16 hrs of Nx or Hx followed by 3 hrs of LPS (100 ng/mL); after 3 hrs, we isolated total RNA for sequencing analyses. As expected, LPS alone in Nx elicited a global change in gene expression, upregulating 4,047 genes (Figure 2A; left). In comparison, Hx alone upregulated 1,119 genes. There were 427 overlapping genes that were upregulated both by Hx and by LPS (Figure 2B; top), demonstrating that LPS has additional effects on global gene transcription that are unique from Hx. Importantly, Hx pre-exposure prior to LPS addition enhanced the expression of 2,313 genes (Figure 2A; right). Of these genes, 698 were a consequence of Hx alone, leaving 1,615 genes that were enhanced by LPS with Hx without being programmatically increased by Hx alone (Figure 2B; Bottom). These are genes that we refer to as “primed” because their gene expression was unchanged by the Hx stimulus alone but was still enhanced by Hx exposure prior to LPS stimulation (when compared to LPS stimulation alone).

**Figure 2.**
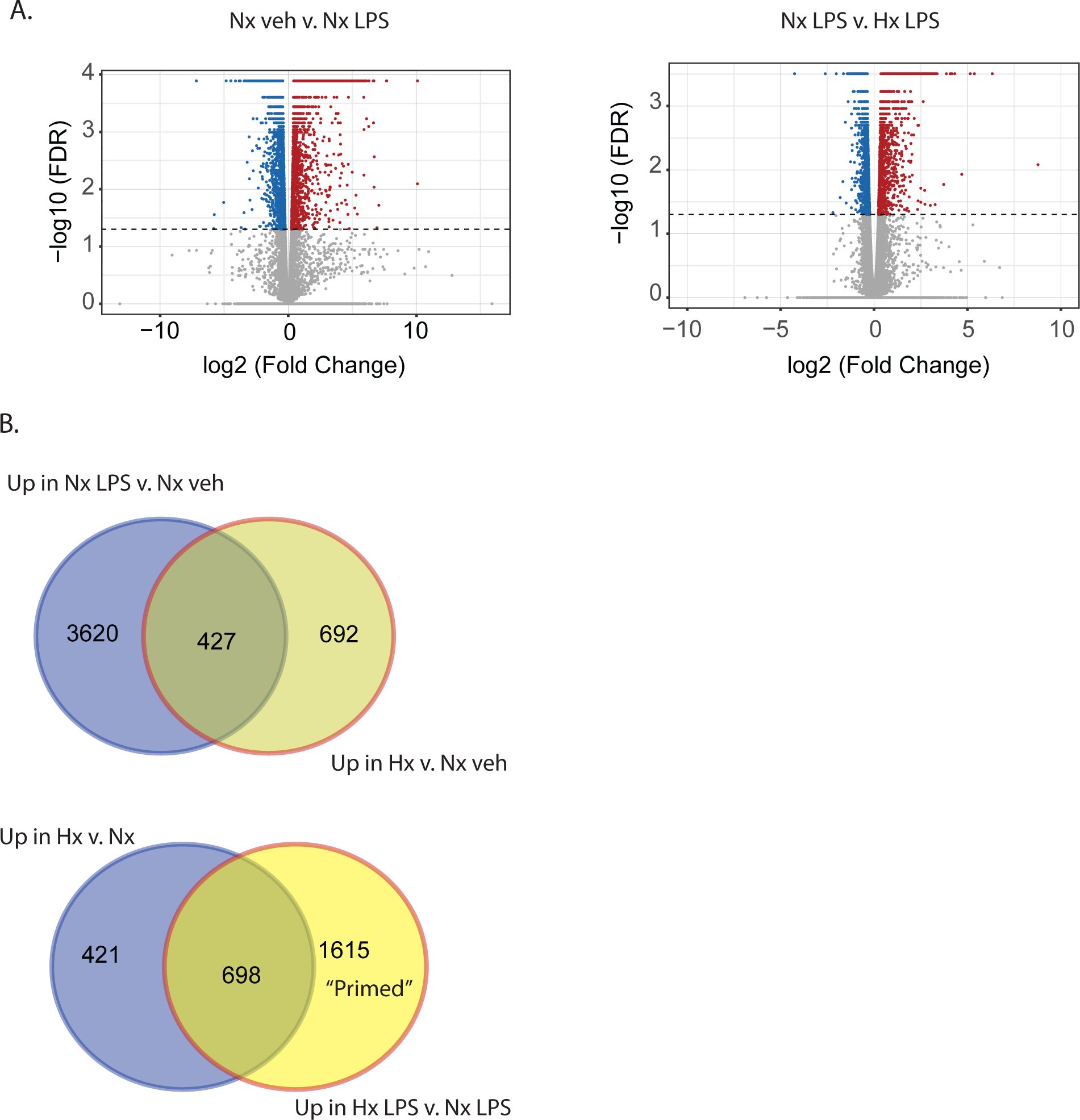
Hypoxia primes LPS-induced gene expression. A) Volcano plots of differential gene expression changes in microglia exposed to Nx LPS vs Nx veh (left) or Nx LPS vs. Hx LPS (right). Results were analyzed using CummeRbund. Red dots represent significantly (FDR < 0.01) upregulated genes and blue dots represent significantly downregulated genes. B) Venn diagrams comparing the number of unique and similar genes upregulated in Nx LPS vs. Nx to genes upregulated by Hx (top) vs. Nx and venn diagrams comparing the number of unique genes upregulated in Hx vs. Nx compared to genes upregulated by Hx LPS vs. Nx LPS. There are 1,615 “primed” genes that are enhanced in Hx LPS vs. Nx LPS that are not upregulated by Hx alone. Results are derived from n = 3 biological replicates/treatment.

We next used KEGG Pathway analysis to categorize the types of genes upregulated by LPS and those primed by Hx (Figure 3). We specifically examined the top 10 most significantly enriched gene categories. As expected, many LPS-induced genes were associated with immune-related categories such as “cytokine-cytokine receptor interactions” and “complement and coagulation cascades”, and infections such as “Influenza A”. Many of the gene categories identified following LPS treatment were also enriched in the primed genes (red bars indicate gene categories shared with LPS-induced genes).

**Figure 3.**
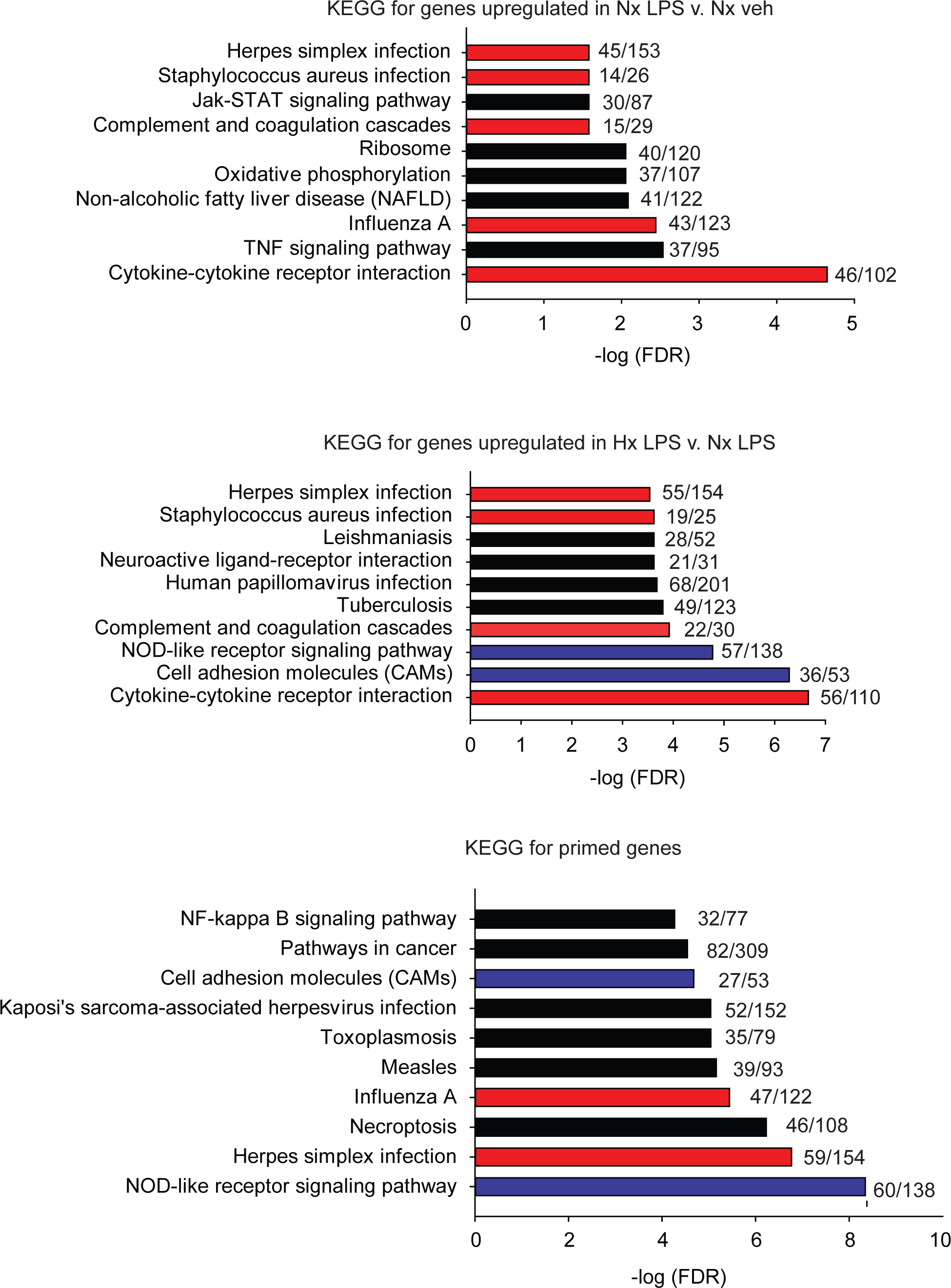
KEGG pathway analyses for all significantly upregulated genes in Nx LPS vs. Nx, Hx LPS vs. Nx LPS, and Primed genes. Each graph displays the top 10 most significant categories. Numbers at the end of each bar are the number of observed/expected genes in each category. Red bars represent LPS-induced gene categories that are shared with Hx-induced and primed gene categories. Blue bars represent Hx-induced gene categories shared with primed gene categories. Black bars represent all unshared categories. Gene expression results are derived from n = 3 biological replicates/treatment.

However, many gene categories were also unique to Hx, such as “cell adhesion molecules” and “NOD-like receptor signaling pathway” (categories represented by blue bar are enriched in Hx conditions only). Importantly, Hx-primed genes, genes that were not augmented by Hx alone but whose expression was enhanced in the presence of LPS, had unique categories that included “pathways in cancer” and “NF-ĸB signaling pathway”. These results suggest that Hx-primed genes may be unique players in inflammatory processes.

Lastly, we identified which Hx primed genes had the largest fold change in expression following LPS treatment. We took the top 100 primed genes with the largest fold changes and submitted them to STRING analyses to identify relationships between the proteins encoded by those genes (Figure 4). For these analyses, we used the “highest confidence” settings and removed unconnected nodes. The results demonstrated that IL-1β and JAK2 were two of the nodes with the most connections, and the genes encoding IL-1β, IL-1α, and NOS2 had some of the largest fold changes in expression. These findings suggest that the largest Hx primed gene changes occur in those encoding pro-inflammatory molecules.

**Figure 4.**
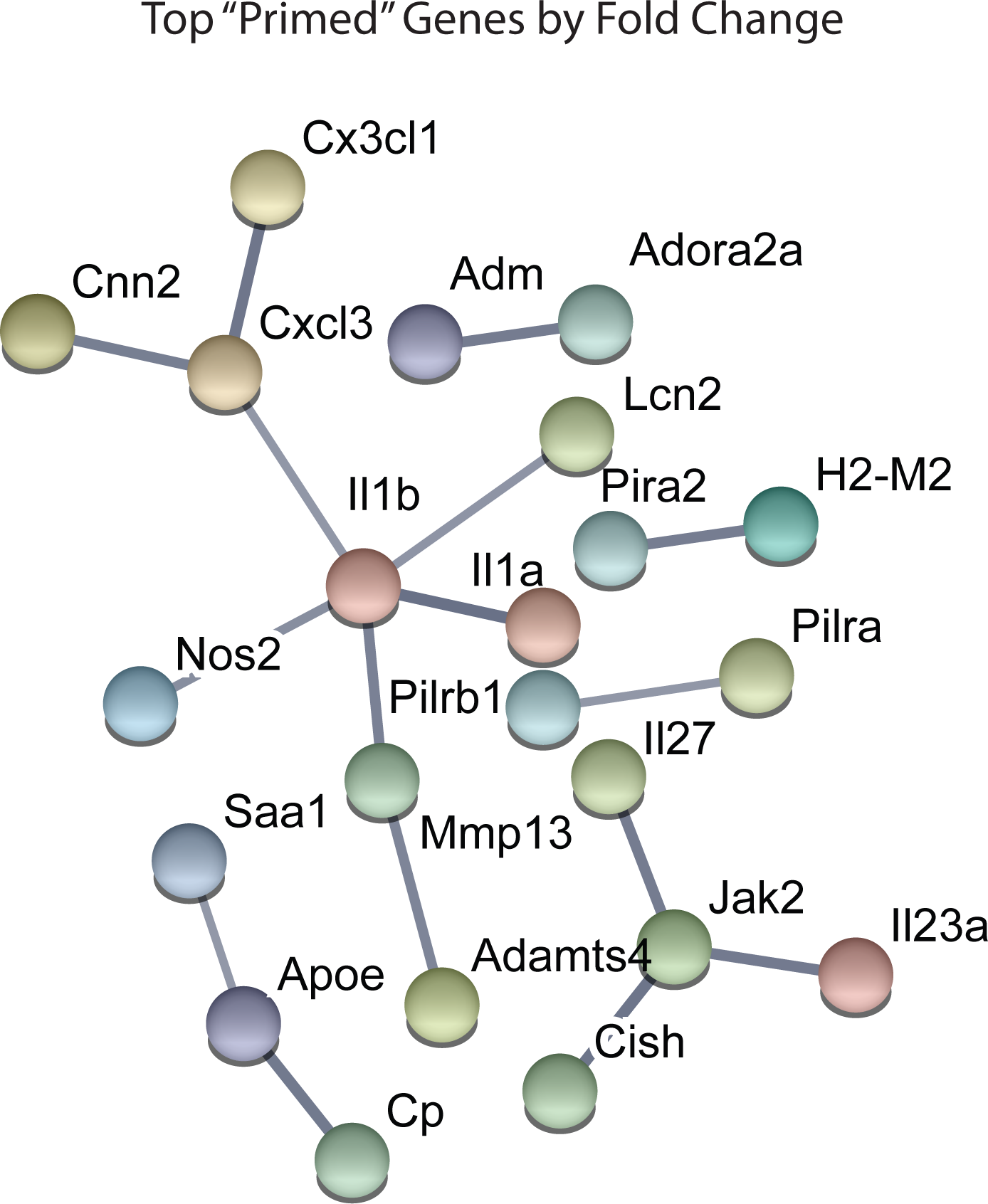
STRING network analyses reveal primed genes are pro-inflammatory. Analyses were performed on the top 100 primed genes with the largest fold changes. Gene expression results are derived from n = 3 biological replicates/treatment.

### Hx induces global enrichment of H3K4me3 at immune-and glycolysis-related genes, but not at primed genes

Previous studies examining HIF-1α-dependent gene priming demonstrate that H3K4me3 is enriched at primed genes (20, 21, 29), possibly accounting for long-term enhancement of subsequent inflammatory responses. Thus, we tested if Hx-primed genes similarly displayed H3K4me3 enrichment at gene promoters in microglia. Using H3K4Me3 ChIP-seq followed by peak analyses in Homer (42), we identified 17,658 unique H3K4me3 peaks that were induced by Hx (Figure 5A). As expected, most of these peaks fell near transcription start sites (TSS; Figure 5B). While many of these peaks were found at the TSS of glycolysis-related genes like *Vegfa*, pro-inflammatory cytokines like *Il1b* did not have peaks enriched at the TSS (Figure 5C). *Il1a* also did not have a peak at the TSS, but it should be noted that *Il1a* has a non-canonical promoter lacking TATA and CAAT box regulatory regions (45) and that *Il1a* expression is regulated by a downstream anti-sense transcript in the region demarcated Gm14023 (46). This downstream genomic feature did display an Hx-induced peak (Figure 5C).

**Figure 5.**
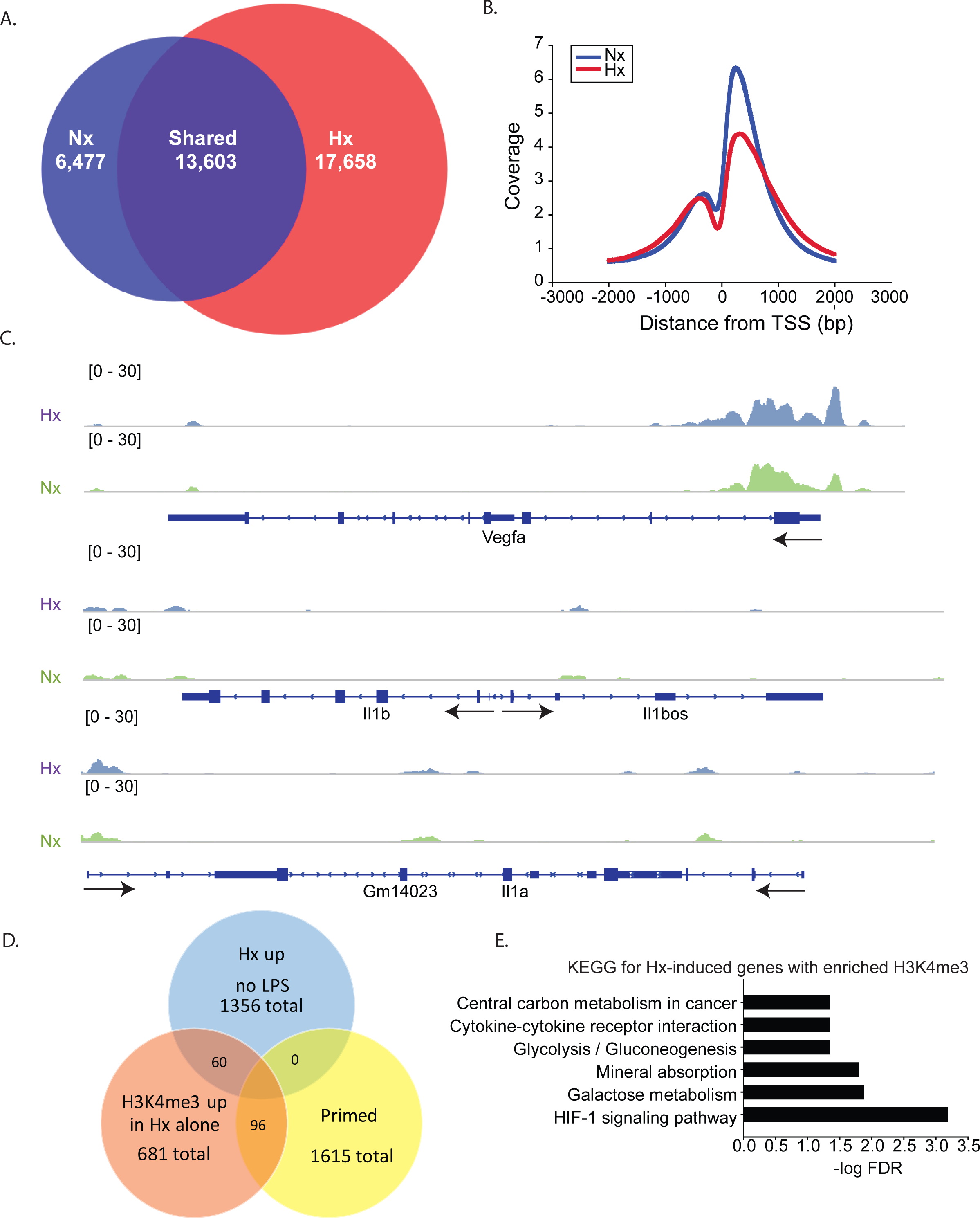
Hypoxia increases global H3K4me3, but not at primed genes. A) Venn diagrams showing the total number of all unique and shared H3K4me3 peaks enriched 4-fold above 10% input in Nx and Hx conditions. Peak analyses were performed on merged consensus tag directories in n = 3 replicates/treatment. B) Histogram of the location of all H3K4me3 peaks enriched in Nx (blue line) and Hx (red line) relative to the transcription start site of each gene. Coverage is defined as the ChIP fragment depth (per bp per peak). C) Representative IGV images of H3K4me3 at VEGF, a HIF-1 target gene, and at Il1b and Il1a, two of the top 100 primed genes with the largest fold changes in gene expression. D) Venn diagram showing the number of Hx-induced genes vs. “primed” genes that have increased H3K4me3 at the transcription start site after Hx exposure. E) KEGG Pathway analyses of the 60 genes upregulated by Hx that also have concomitant increase in H3K4me3 at the transcription start site.

When examining the H3K4me3 peaks that were increased by Hx and that fell within -1kb and +300 bp of the gene TSS, we identified only 96 peaks correlating with genes that were primed by Hx (Figure 5D and Supplemental Figure 2). These observations suggest that most primed genes are not associated with Hx-induced H3K4me3 enrichment, unlike what has been reported by others (20, 29, 47). However, it should be noted that our analyses only examined peaks that were increased by Hx, and that many primed genes already possessed H3K4me3 at the TSS. Nevertheless, we conclude that increases in H3K4me3 do not likely account for gene priming effects induced by Hx. We next examined genes that were upregulated by Hx and that also had concomitant increases in H3K4me3 (Figure 5E), genes that were not “primed” by definition. KEGG Pathway analyses confirmed that these genes were most enriched in the categories of “HIF-1 signaling pathway” as well as “cytokine-cytokine receptor interaction”, indicating that Hx, as expected, enriches H3K4me3 at genes upregulated by Hx, which are involved in immune- and glycolysis-related pathways.

### Hypoxia and β-glucan increase H3K4me3 at unique genes involved in cell metabolism

Given that our results support an H3K4me3-independent mechanism of gene priming, we compared our H3K4me3-induced peaks to those identified in human monocytes primed with β-glucan ((20); Figure 6). The β-glucan dataset had much greater H3K4me3 enrichment than the Hx dataset. Therefore, we analyzed the β-glucan dataset using two peak thresholds: 1.4-fold above background (Figure 6) and 4-fold above background (Supplemental Figure 3). We compared the peaks identified using both thresholds to those we identified in our Hx dataset (which used the 1.4-fold threshold); both comparisons yielded similar outcomes in terms of the gene categories enriched with H3K4me3. When peaks were called at 1.4-fold above background, β-glucan induced ∼8000 H3K4me3 peaks at gene promoters, whereas Hx only induced ∼700 H3K4me3 peaks (Figure 6A). Interestingly, more than 50% of Hx-induced H3K4me3 overlapped with β-glucan-induced peaks (Figure 6A). We analyzed this list of overlapping genes using STRING (44). While no KEGG Pathway categories were enriched with our Hx gene list, gene ontology for Biological Process revealed that H3K4me3 peaks shared between the Hx and β-glucan datasets occurred at genes involved in metabolic processes (Figure 6B). Together, these results suggest that H3K4me3 enrichment occurs at metabolic genes, rather than immune-related genes.

**Figure 6.**
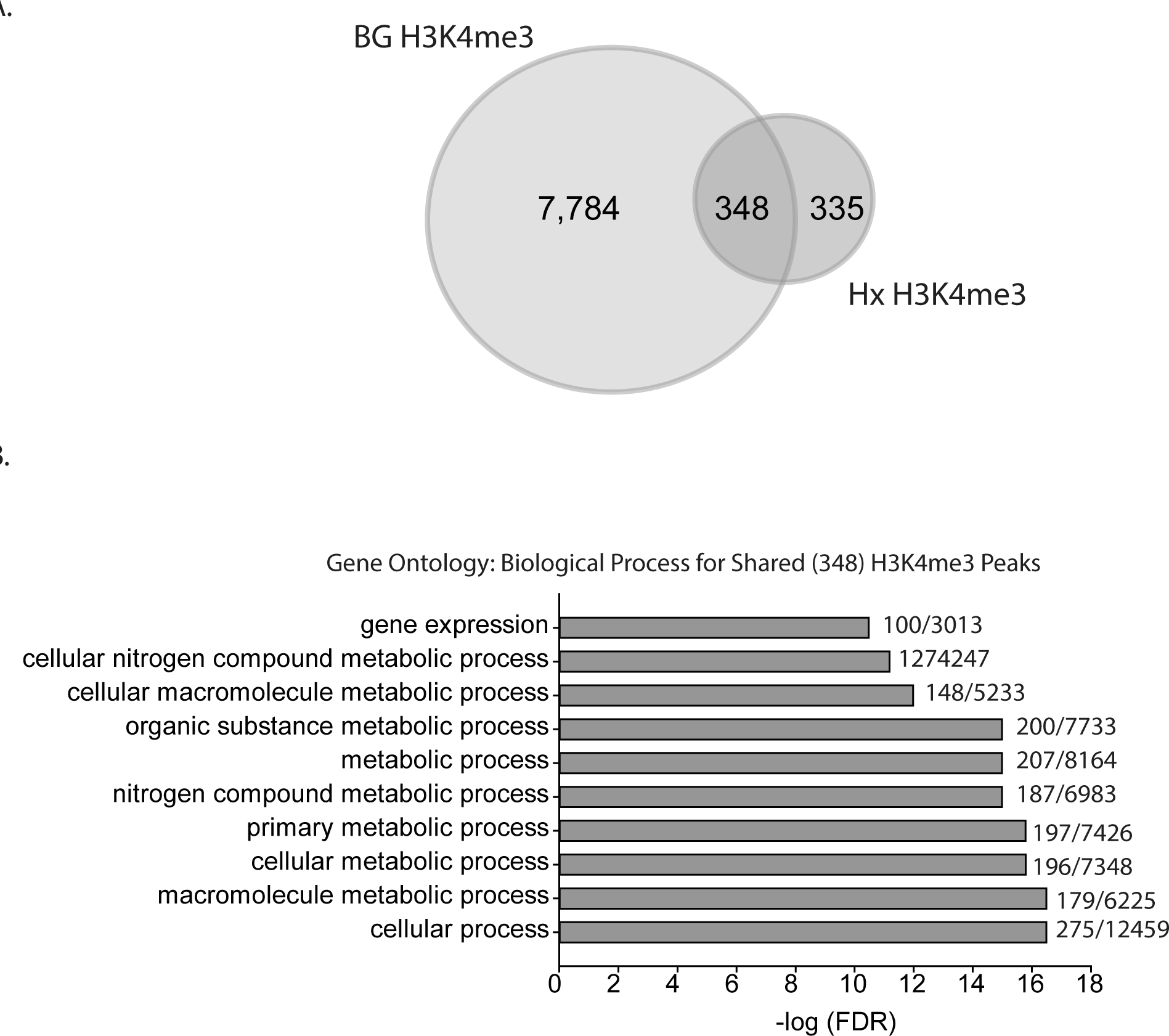
Hypoxia-induced H3K4me3 enrichment occurs in distinct genes from those primed by beta glucan. A) Venn diagram comparing numbers of similar and unique genes with H3K4me3 enrichment after BG or Hx priming. B) Gene Ontology for Biological Process was performed for genes enriched with H3K4me3 at the transcription start site shared between Hx and BG priming. H3K4me3 peaks were called using Nx control or RPMI control as background for Hx and BG priming, respectively. Numbers at the end of each bar represent observed genes/expected genes in each category. Results are derived from n = 3 biological replicates/treatment.

### MAGICTRICKS reveals multiple transcription factors that putatively regulate Hx primed genes

Since changes in H3K4me3 do not likely account for Hx-induced gene priming, we used MAGICTRICKS (40) to explore other putative chromatin modifiers that could be associated with Hx-induced gene priming. MAGICTRICKS utilizes a matrix of maximum ChIP-seq scores for every transcription factor at every gene analyzed in the ENCODE database (48). We added to this matrix a HIF-1α track (GSM3417785_16h_PM14_1_peaks from GEO # GSM3417785) from cells exposed to similar paradigms of Hx described here, and we confirmed that this track recognized HIF-1α regulated genes by analyzing our list of genes upregulated by Hx (Supplemental Figure 4). We then used MAGICTRICKS to analyze our list of genes primed by Hx (Figure 7). Results showed that our gene list was significantly enriched for multiple transcription factors and cofactors related to immune system function including Stat3, Ezh2, and NF-ĸB (Figure 7A, B) and surprisingly, not HIF-1α. These results indicate that mechanisms of gene “priming” induced by Hx are distinct from those genes that are simply upregulated by Hx alone, and that there are likely multiple factors regulating the expression of primed genes.

**Figure 7.**
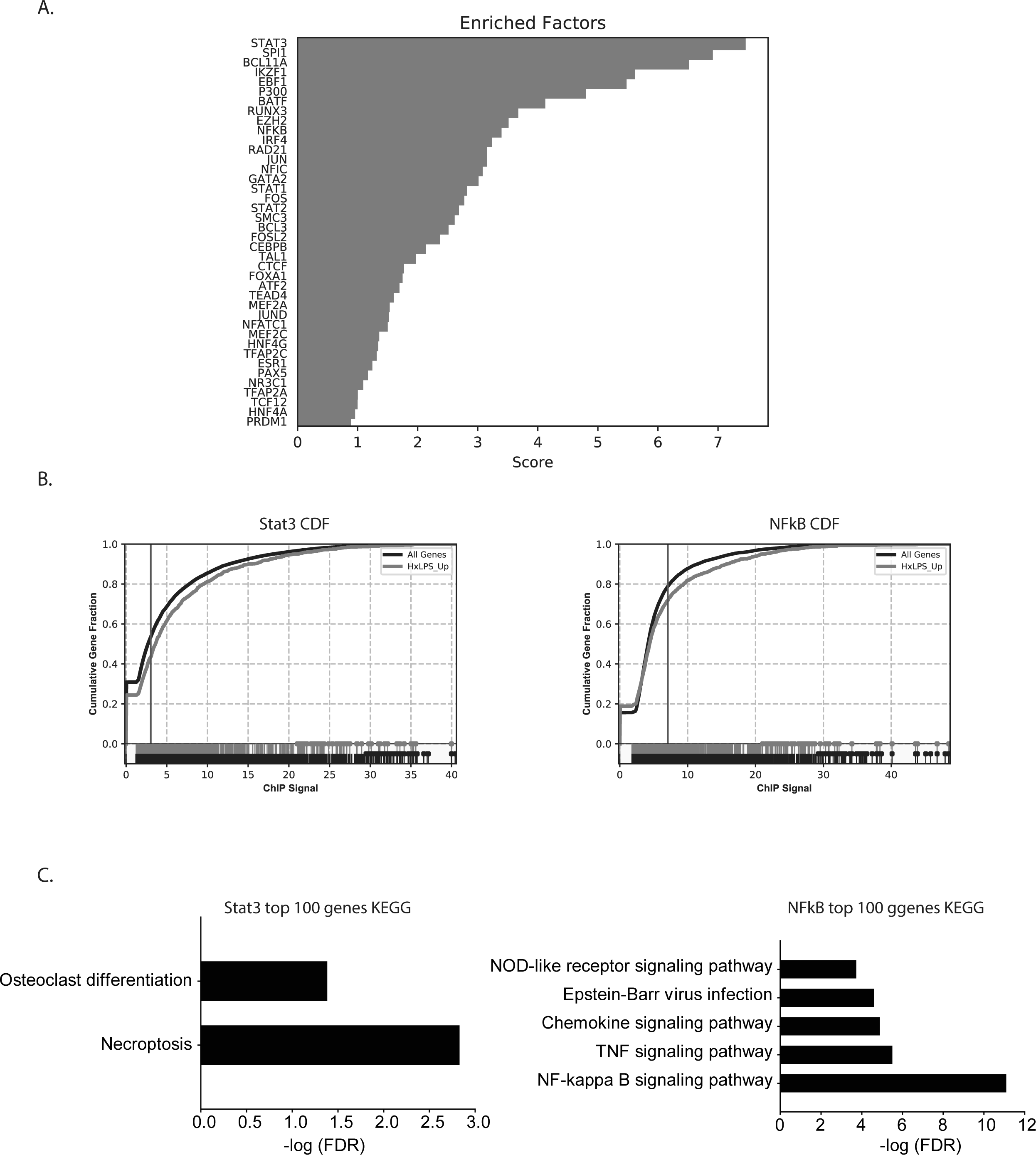
MAGICTRICKS analyses reveal multiple putative transcription factors that bind at “primed” genes. A) MAGICTRICK summary output for Hx primed genes. B) Cumulative distribution functions for the transcription factors Stat3 (left) and NF-ĸB (right). C) Target genes for Stat3 or NFĸB that were identified using MAGICTRICKS were sorted by ChIP score (highest to lowest), and the top 100 genes were examined using KEGG Pathway analyses. Different KEGG categories were enriched with Stat3 genes (left) vs. NFĸB genes (right). Results are derived from n = 3 biological replicates/treatment.

Since KEGG and STRING analyses had previously identified the NF-ĸB pathway as a category of genes primed by Hx (Figures 3 and 4), we compared the MAGICTRICKS list of genes regulated by Stat3 vs. NF-ĸB; Stat3 was the top transcription factor identified by MAGICTRICKS as the most common factor involved in regulating the Hx-primed genes. Thus, we took the genes with the top 100 ChIP scores for NF-ĸB and Stat3 and subjected them to KEGG Pathway analysis (Figure 7C). Interestingly, for Stat3, only two categories were enriched, “osteoclast differentiation” and “necroptosis”. However, for NF-ĸB, multiple immune signaling-related categories were enriched including NF-ĸB pathway. These results reveal very different functions for the genes that are regulated by Stat3 vs. NF-ĸB, and they demonstrate that the priming of pro-inflammatory cytokines may be more regulated by NF-ĸB.

### Pharmacological NF-ĸB inhibition attenuates Hx-induced inflammatory gene priming

The MAGICTRICKS results suggested that NF-ĸB may be a mediator of Hx-primed pro-inflammatory genes. Thus, to test if NF-ĸB signaling contributed to Hx gene priming, we cultured microglia in Hx or Nx in the presence or absence of BOT-64, a reversible inhibitor of I kappa kinase 2 (IKK2), which prevents NF-κB activation (Figure 8). We then washed out the inhibitor, stimulated with LPS, and isolated cells for gene expression analyses of *Il1a* and *Il1b*, two of the cytokines with the largest fold gene changes in LPS-induced transcriptome analyses after Hx priming (Figure 4).

**Figure 8.**
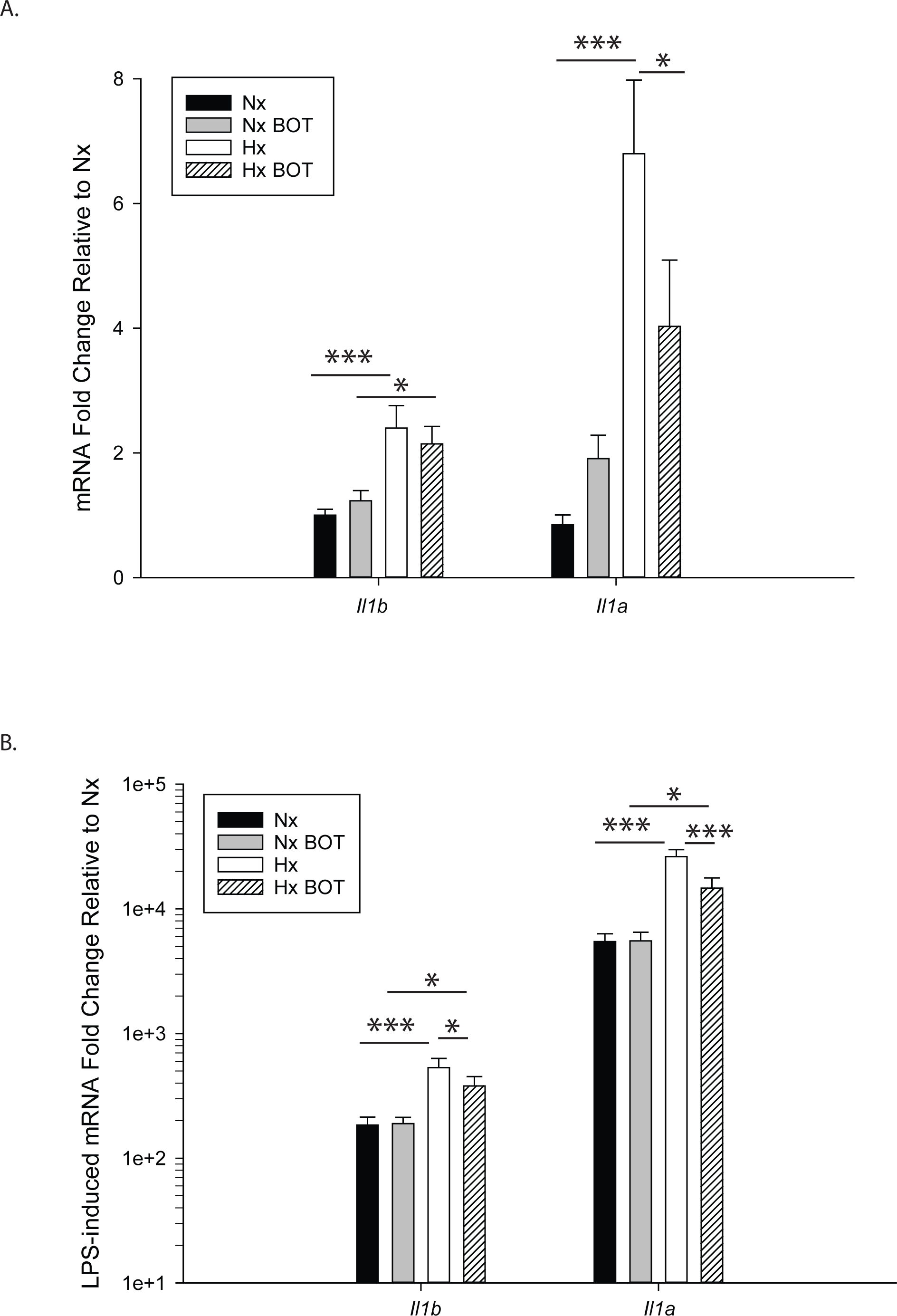
Pharmacologic NFĸB inhibition attenuates hypoxia-induced inflammatory Il-1β and Il-1α gene priming. N9 microglia were exposed to Nx or Hx in the presence of vehicle or the IKK2 inhibitor BOT-64 (3 µM) after which time the inhibitor was washed out. Fresh medium was replaced, and the cells were challenged with LPS (100 ng/ml) for 3 hrs. Inflammatory gene expression was evaluated by qRT-PCR. (A) Il-1β and Il-1α gene expression was evaluated in the presence of BOT-64 treatment during Hx. (B) LPS-induced Il-1β and Il-1α gene priming was evaluated in the presence of BOT-64 treatment during Hx. Results are presented as the average fold change ± SEM expressed relative to Nx vehicle, of n = 5 independent experiments. All results are displayed as the average fold change ± SEM expressed relative to WT Nx vehicle of n =6-7 independent experiments. *p < 0.05, **p < 0.01, ***p < 0.001; Two-way RM-ANOVA.

Results showed that in vehicle-treated control cells, there was no main effect of BOT on *Il1b* (p=0.932) nor *Il1a* (p=0.200) mRNA levels (Figure 8A; Table 3)). However, there was a main effect of Hx on *Il1b* (p= 0.003) and *Il1a* (p=0.003) gene expression. Posthoc analyses revealed that Hx had a small, but statistically significant effect on *Il1b* (p< 0.001) and *Il1a* (p < 0.001), increasing their expression approximately 2-fold and 6-fold, respectively, relative to Nx vehicle conditions. Interestingly, in the presence of NF-ĸB inhibition, Hx lost its ability to upregulate *Il1a* gene expression relative to Nx (p = 0.104), suggesting that NF-ĸB contributes to Hx-induced upregulation of *Il1a* expression. Following LPS treatment (Table 4; Figure 8B), there was a main effect of Hx on both *Il1b* (p=0.005) and *Il1a* (p=0.003) expression. Posthoc analyses revealed that Hx significantly enhanced (primed) LPS-stimulated *Il1b* expression by about 2-fold (p< 0.001) and enhanced *Il1a* expression by about 4-fold (p < 0.001). There was a main effect of IKK2 inhibition on *Il1a* expression (p=0.003), but not on *Il1b* expression (p=0.108; Figure 8B). Additionally, there was a significant interaction between IKK2 inhibition and Hx for *Il1a* (p= 0.020), and a trending interaction for *Il1b* expression (p=0.085). Posthoc analyses showed that BOT treatment and washout prior to LPS exposure was effective as it did not significantly alter LPS-induced levels of *Il1b* (p=0.929) nor *Il1a* (p=0.959) in Nx. Importantly, after BOT exposure during Hx, there was a significant reduction in the magnitude of Hx-induced gene enhancement by LPS, both for *Il1b* (p = 0.026) and *Il1a* (p = 0.016) gene expression. BOT reduced Hx gene priming of *Il1b* by approximately 50% (p= 0.016) and *Il1a* by 60% (p< 0.001). Overall, these data support the idea that NF-ĸB contributes to hypoxia-induced priming of pro-inflammatory cytokines.

**Table 3.** Two-way ANOVA Results for NF-kB Inhibition without LPS

| Gene | n/tx (Nx veh, Nx BOT, Hx veh, Hx BOT) | Main Effect of Hx |  | Main effect of BOT |  | Interaction Hx v. BOT |  |
| --- | --- | --- | --- | --- | --- | --- | --- |
|  |  | F | P | F | P | F | P |
| *Il1b* | 7,7,7,7 | 21.668 | 0.003 | 0.00783 | 0.932 | 1.641 | 0.247 |
| *Il1a* | 6,6,6,6 | 24.581 | 0.003 | 2.074 | 0.200 | 8.066 | 0.047 |

**Table 4.** Two-way ANOVA Results for NF-kB Inhibition with LPS

| Gene | n/tx (Nx LPS, Nx<br>BOT LPS, Hx LPS,<br>Hx BOT LPS) | Main Effect of Hx |  | Main effect of BOT |  | Interaction Hx v.<br>BOT |  |
| --- | --- | --- | --- | --- | --- | --- | --- |
|  |  | F | P | F | P | F | P |
| *Il1b* | 7,7,7,7 | 18.850 | 0.005 | 3.564 | 0.108 | 4.243 | 0.085 |
| *Il1a* | 6,6,6,6 | 28.580 | 0.003 | 20.936 | 0.003 | 13.828 | 0.020 |

### Hypoxia primes microglial pro-inflammatory gene expression 3 and 6 days after exposure

We lastly tested if Hx could enhance microglial responses to inflammatory stimuli multiple days after the removal of the initial Hx exposure. For these experiments, we used primary microglia because they can be cultured for multiple days without the need to passage. Remarkably, we found that 3 days (Figure 9A; Table 5) and 6 days (Figure 9B) post-Hx, microglia still exhibited potentiated LPS-induced *Il1a* and *Il1b* gene responses. At 3 days post-Hx, there was a significant interaction between Hx and LPS exposure for *Il1b* (p = 0.038) and *Il1a* (p = 0.014). Posthoc analyses confirmed that LPS-induced *Il1b* gene expression was 2-fold higher in microglia pre-exposed to Hx compared to microglia previously exposed to Nx (p = 0.002). Similarly, LPS-induced *Il1a* gene expression after Hx exposure was ∼2.2 fold higher than after Nx exposure (p < 0.001). At 6 days post-Hx (Figure 9B; Table 6), a primed *Il1a* response to LPS (p =0.013) and a trending *Il1b* response (p = 0.094) persisted, suggesting that Hx leads to modest (20-50%), yet long-lasting enhancement of LPS inflammatory gene responses.

**Figure 9.**
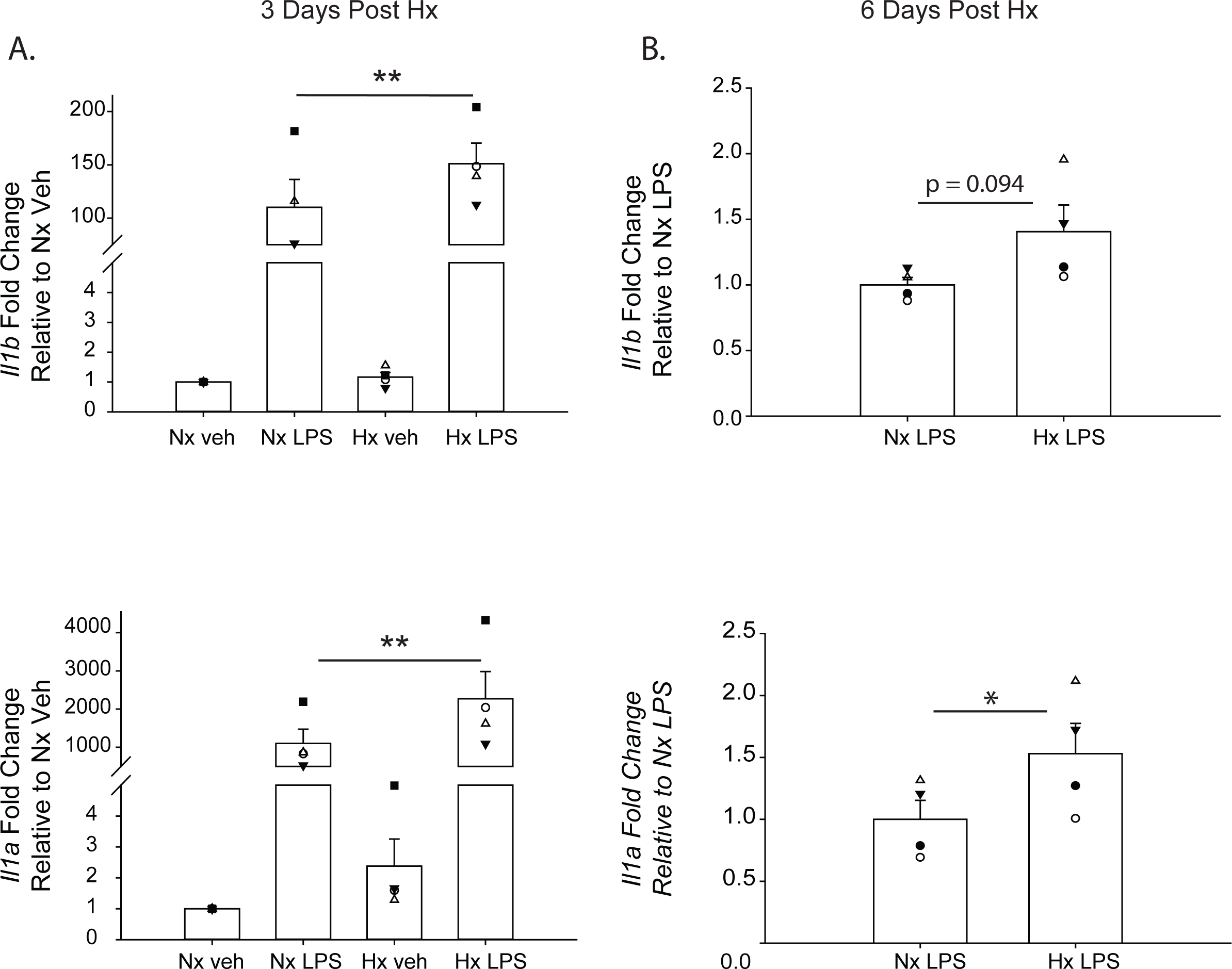
Microglia exhibit gene-specific memory of hypoxia exposure that lasts for several days. Primary microglia were cultured overnight in Nx or Hx. The cells were washed with fresh medium containing MCSF, and then cultured for 3 (A) or 6 (B) days in Nx. Cells were then treated with LPS (100 ng/ml) for 3 hrs and harvested for qRT-PCR analysis of inflammatory gene expression. (A) Cells were cultured for 3 days post-Hx, treated with LPS, and harvested for qRT-PCR analysis of Il1b (top) and Il1a (bottom). Values are expressed as the average fold change ± SEM relative to Nx vehicle of n=4 independent experiments. * p < 0.05, **p < 0.01 vs. Nx LPS; Two-way RM ANOVA. (B) Cells were cultured for 6 days post-Hx. Values are expressed as the average fold change ± SEM relative to Nx LPS of n=4 independent experiments. *p < 0.05 vs. Nx LPS; paired T-test.

**Table 5.** Two-way ANOVA Results for Hx Priming 3-Days Post-Hx

| Gene | n/tx (Nx veh, Nx<br>LPS, Hx veh, Hx<br>LPS | Main Effect of Hx |  | Main effect of LPS |  | Interaction Hx v.<br>LPS |  |
| --- | --- | --- | --- | --- | --- | --- | --- |
|  |  | F | P | F | P | F | P |
| *Il1b* | 4,4,4,4 | 13.187 | 0.036 | 27.248 | 0.014 | 12.736 | 0.038 |
| *Il1a* | 4,4,4,4 | 27.100 | 0.014 | 21.569 | 0.019 | 27.301 | 0.014 |

**Table 6.** Paired-T-test Results for Hx Priming 6 Days Post-Hx

| Gene | n/tx (Nx LPS, Hx LPS) | Effect of Hx |  |
| --- | --- | --- | --- |
|  |  | t | P |
| *Il1b* | 4,4 | -2.416 | 0.094 |
| *Il1a* | 4,4 | -5.249 | 0.013 |

## DISCUSSION

We tested the overall hypothesis that Hx, like low dose pathogen exposure, can prime the microglial inflammatory gene response to a subsequent inflammatory stimulus *in vitro*. Our results demonstrated that a Hx stimulus that activates glycolytic metabolic pathways can robustly prime microglial responses to a subsequent inflammatory challenge, both within 3 hrs and up to 6 days following the removal of Hx. Although these are *in vitro* results, they are consistent with the pathogen priming literature in macrophages that was also done *in vitro*. That Hx augments microglial inflammatory gene expression has significant relevance in the context of Hx brain injuries, including ischemic stroke, in which the CNS tissue becomes completely anoxic (49). Given that multiple Hx brain disorders, including stroke, are associated with neurodegenerative disease (50) and aberrant inflammatory microglial activities (51), the results presented here provide evidence for a novel mechanism whereby microglia can be altered long-term by hypoxic brain injury, potentially making the brain more susceptible to a subsequent or ongoing injury or disease.

While our results indicate that Hx has profound effects on microglial inflammatory activities, the level of Hx and the cellular microenvironment are likely important determinants of any long-term effects of Hx pre-exposure, particularly *in vivo*. Other studies have demonstrated that certain paradigms of sustained Hx *in vivo* can actually dampen peripheral leukocyte glycolytic metabolism long-term (52), leading to improved survival outcomes after subsequent pathogen infection. Although this is consistent with our findings that Hx can have long-lasting effects on microglial inflammatory gene expression, they also contrast with our observations that Hx enhances genes related to glycolytic metabolism. Additional evidence signifying the importance of the Hx paradigm on microglial functions, neonatal microglia exposed to intermittent Hx do not exhibit upregulated glycolysis-related gene expression and further, have attenuated pro-inflammatory responses acutely following intermittent Hx exposure (53).

Therefore, the baseline changes we observed as a result of Hx alone, including increases in glycolysis-related gene expression, may be an important indication of whether microglia will exhibit enhanced or attenuated gene responses to a subsequent inflammatory stimulus.

The idea that cellular metabolism plays a critical role in Hx priming effects is supported by the similarities between our Hx priming in microglia and pathogen priming in peripheral monocytes. Both models demonstrate increased expression of immune- and glycolysis-related genes, and concomitant H3K4me3 enrichment at many of these genes (20, 21, 29, 54). However, unlike the β-glucan and pathogen priming literature (20, 47), we find that the majority of inflammation (i.e. LPS)-induced genes that are enhanced by Hx pre-exposure, including pro-inflammatory cytokines, are not further enriched with H3K4me3 following Hx exposure. This suggests that alternative mechanisms to H3K4me3 at pro-inflammatory genes may play a role in the ability of pro-inflammatory cytokines to be enhanced by subsequent stimuli long-term. Since the most significant categories of genes enriched with H3K4me3 after Hx exposure (including genes shared with the β-glucan dataset) involved cell metabolism, it is likely that H3K4me3 peaks do contribute to long-term Hx effects by changing gene expression related to cell metabolism, thereby promoting a more pro-inflammatory cell phenotype. Additionally, while the evidence presented here suggests that H3K4me3 does not directly regulate pro-inflammatory cytokine gene expression for the majority of primed genes, it does not exclude the possibility that the few pro-inflammatory cytokines that do have increased H3K4me3 (Supplemental Figure 3) are important in the long-term effects of Hx.

In the search for transcription factors and cofactors that could be responsible for regulating Hx-induced gene priming, MAGICTRICKS identified multiple transcription factors including NF-ĸB. Pharmacological studies verified a role for NF-ĸB in primed pro-inflammatory gene regulation, complementing other studies using low-dose LPS as a priming stimulus, which also identified a role for NF-ĸB in enhanced inflammatory gene responses to a subsequent inflammatory challenge (54, 55). However, the results from MAGICTRICKS also revealed that it is very likely that multiple DNA binding factors work in concert with NF-ĸB to regulate Hx-mediated priming of pro-inflammatory genes. Surprisingly, HIF-1α was not a significant regulator of these genes. While HIF-1α likely contributes to gene priming by regulating changes in metabolism-related genes, the MAGICTIRCKS analyses demonstrated that primed genes have a unique set of transcriptional regulators compared to genes that are upregulated by Hx alone.

Altogether, our results demonstrate that *in vitro* Hx primes the microglial inflammatory gene response to a subsequent inflammatory stimulus, both acutely and long-term, in the absence of H3K4me3 enrichment at primed genes. This suggests that Hx is a unique stimulus that when applied alone, does not robustly increase pro-inflammatory cytokine gene expression. However, when it is delivered prior to a more robust inflammatory stimulus, Hx can profoundly enhance the ability of microglia to upregulate pro-inflammatory cytokine expression, thereby potentially enhancing microglia immune function. Thus, Hx pre-exposure may contribute to aberrant CNS processes and provide a mechanism whereby the more complex, long-term effects of Hx can influence microglial function in the context of neural injury and neurodegenerative disease.

## ACKNOWLEDGEMENTS

The authors thank the University of Wisconsin Biotechnology Center DNA Sequencing Facility for providing DNA sequencing facilities and services, the UW Comprehensive Cancer Center Flow Cytometry core (NIH P30 CA014520 and 1S100OD018202-01) and the Bioinformatics Resource Center for help and feedback on data analyses. We also wish to thank Drs. John Svaren and Reid Alisch for helpful discussions and suggestions throughout this project, and for providing guidance and expertise in chromatin immunoprecipitations and ChIP-Seq bioinformatics analyses. Supported by F31NS100229 (EAK) and R01NS085226 (JJW).

