## Supplemental for "Prior hypoxia exposure enhances murine microglial inflammatory gene expression *in vitro* without concomitant H3K4me3 enrichment"

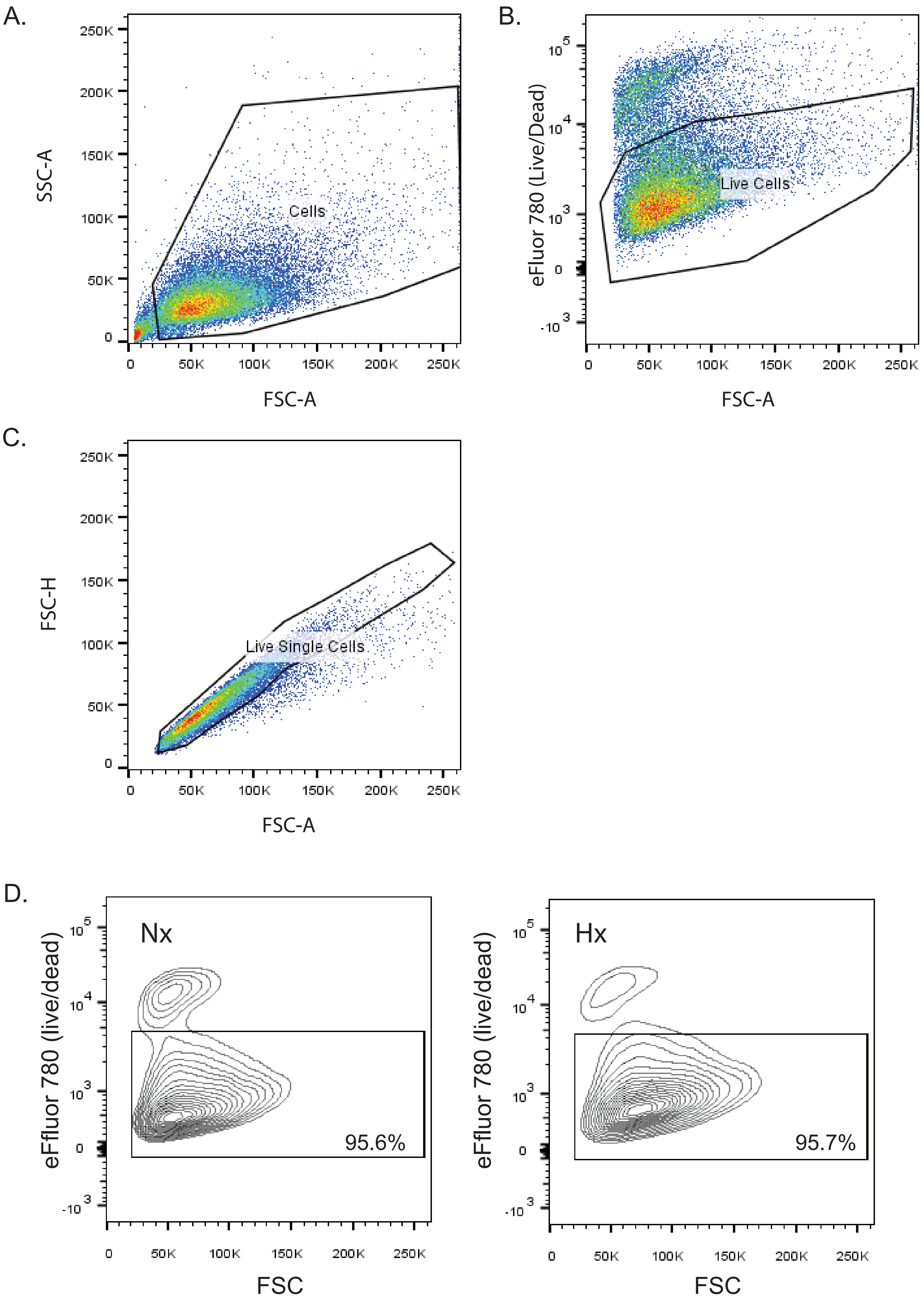
A.
SSC-A
250K
200K
150K
100K
50K
0
Cells
0
50K
100K
150K
200K
250K
FSC-A
B.
eFluor 780 (Live/Dead)
$10^5$
$10^4$
$10^3$
0
$-10^3$
Live Cells
0
50K
100K
150K
200K
250K
FSC-A
C.
FSC-H
250K
200K
150K
100K
50K
0
Live Single Cells
0
50K
100K
150K
200K
250K
FSC-A
D.
Nx
eFfluor 780 (live/dead)
$10^5$
$10^4$
$10^3$
0
$-10^3$
95.6%
0
50K
100K
150K
200K
250K
FSC
Hx
eFfluor 780 (live/dead)
$10^5$
$10^4$
$10^3$
0
$-10^3$
95.7%
0
50K
100K
150K
200K
250K
FSC

**Supplemental Figure 1. Flow cytometry for cell viability.** (A) Cells were distinguished from debris using forward scatter (FSC) and side scatter (SSC). (B) Live cells were gated based on live/dead dye exclusion. (C) Single cells were identified by examining FSC height vs. FSC width. (D) N9 microglia were exposed overnight to 1-1.5% $O_2$ (Hx) or to 21% $O_2$ (room air; Nx). Cells were isolated for flow cytometry and stained with the live/dead dye eFluor780 to assess cell viability.

Kiernan et al.
Supplemental Figure 2

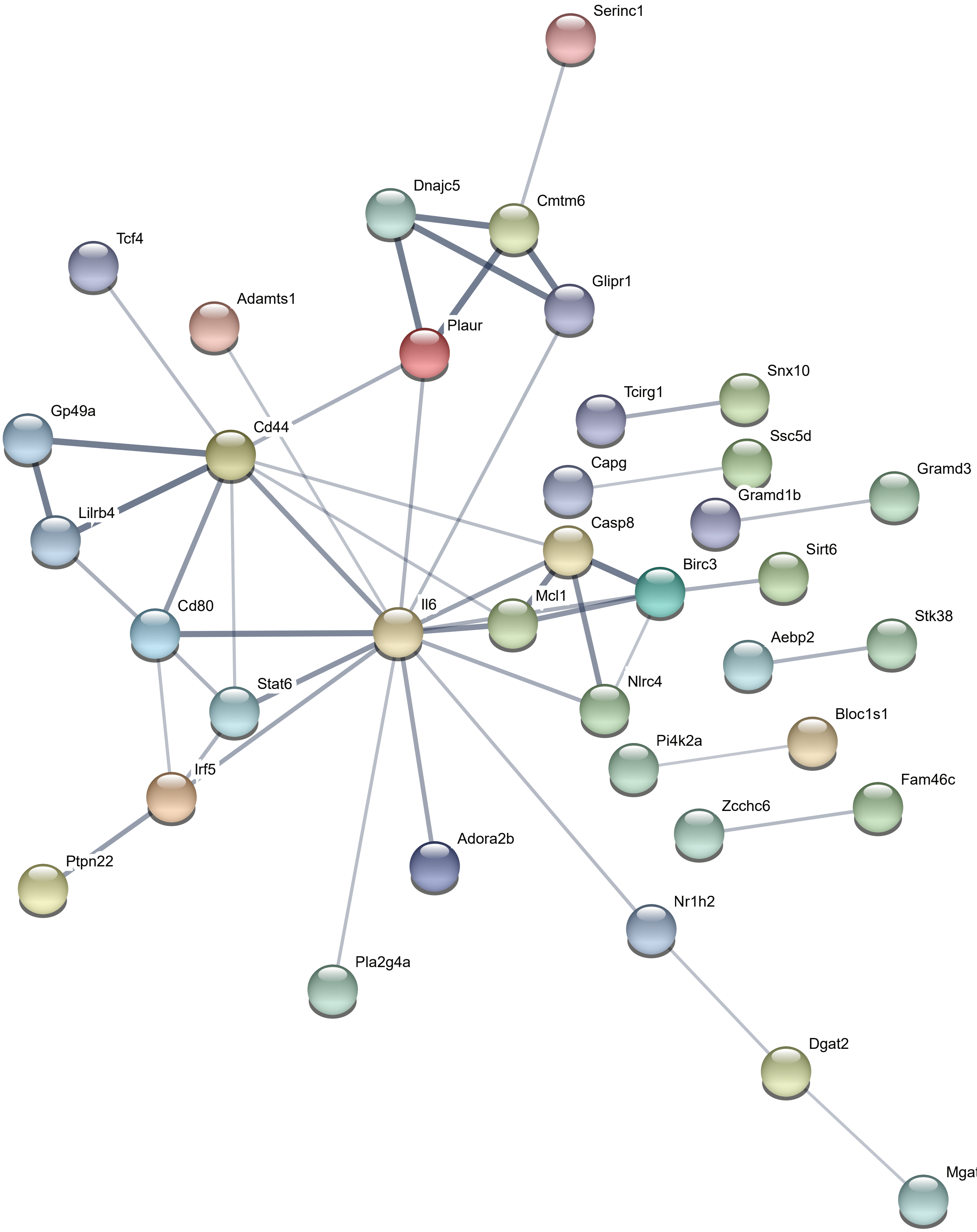

**Supplemental Figure 2. STRING diagram for the 96 primed genes that had concomitant increases in H3K4me3 after Hx exposure.** Setting parameters were changed so that only high confidence connections are shown. All disconnected nodes were removed.

Kiernan et al.
Supplemental Figure 3

A.

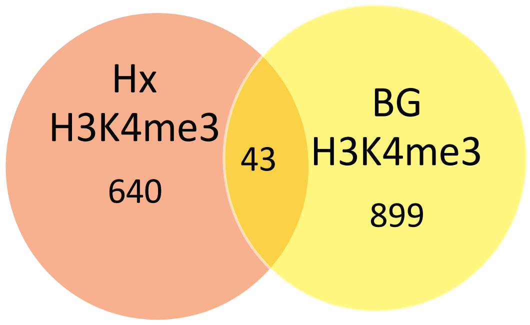


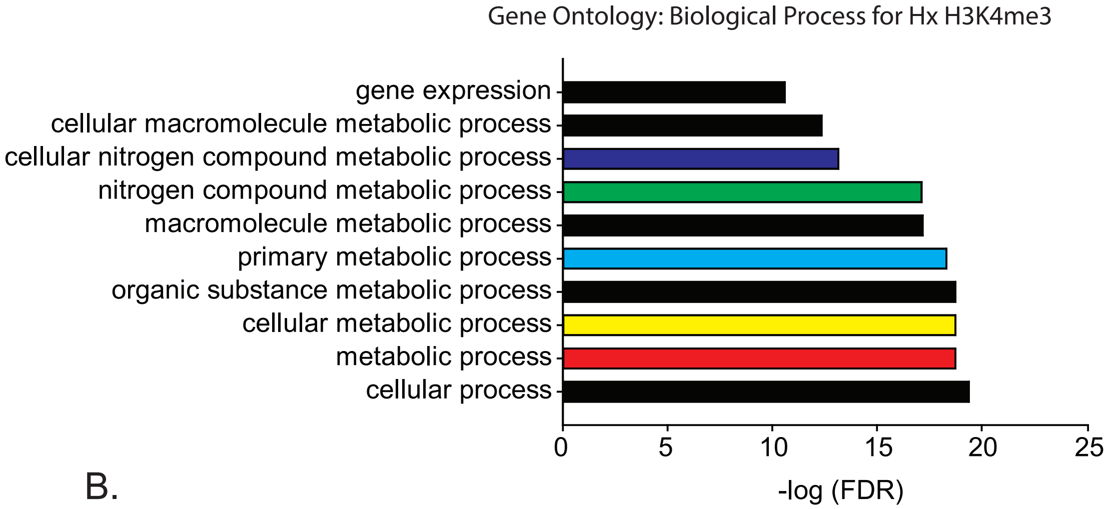


B.

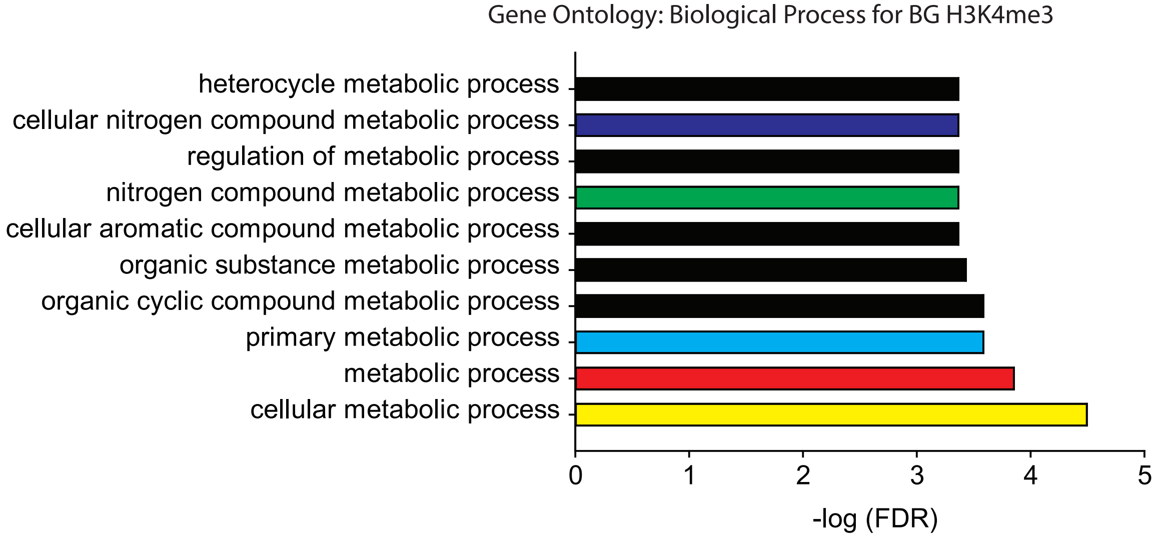

**Supplemental Figure 3. Analysis of H3K4me3 peaks in β-glucan primed human monocytes compared to Hx primed microglia when peaks were called at 4-fold above background.** (A) Venn diagram showing overlap of H3K4me3 peaks in Hx primed vs. β-glucan primed cells. (B) Gene ontology for Biological Process demonstrated that both Hx primed microglia and β-glucan primed monocytes had H3K4me3 peaks at genes involved in cellular metabolism. Shared categories are in color.

MAGIC analysis of genes upregulated by Hx

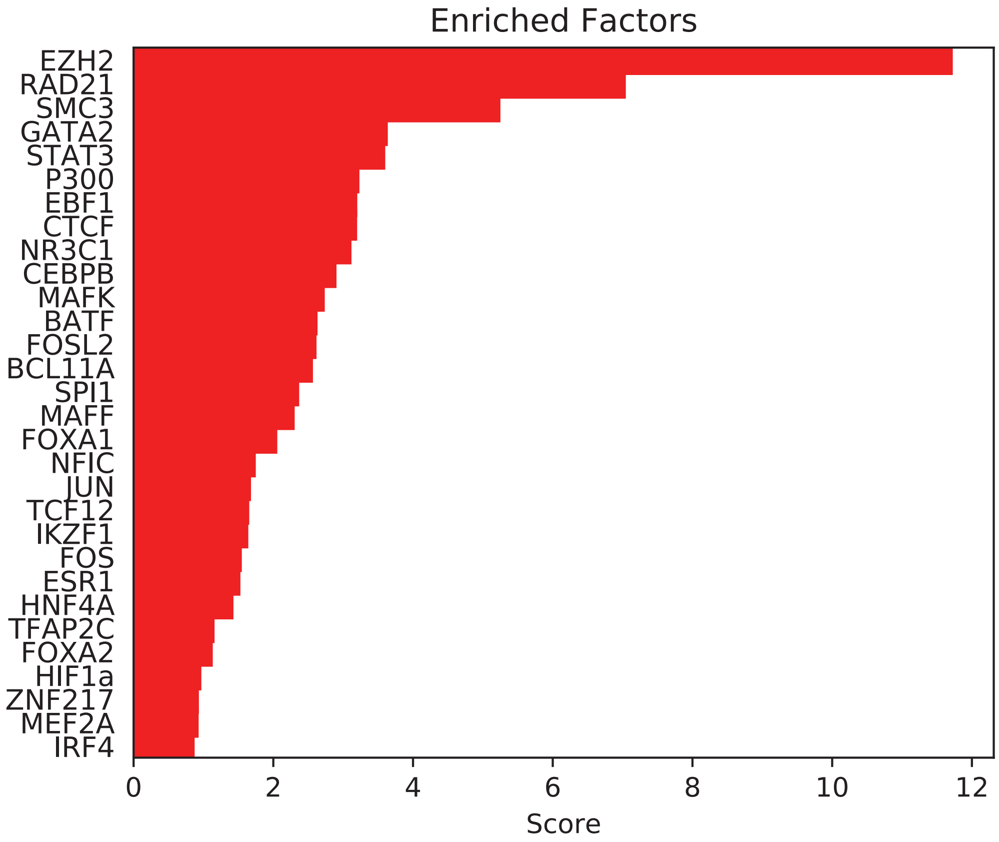

**Supplemental Figure 4. MAGICTRICKS summary score for transcription factors regulating genes upregulated by Hx.** Analyses revealed Hif-1α as a transcription factor that binds genes enriched in the list of Hx upregulated genes.
